# Oxford Nanopore Sequencing as a useful tool for investigating the population dynamics of invasive begomoviruses in Sicily

**DOI:** 10.1101/2025.07.31.667907

**Authors:** Sofia Bertacca, Silvia Rotunno, Fulco Frascati, Emanuela Noris, Gian Paolo Accotto, Salvatore Davino, Laura Miozzi, Anna Maria Vaira

**Affiliations:** Institute for Sustainable Plant Protection, National Research Council, 10135 Torino, Italy; Department of Agricultural, Food and Forest Sciences (SAAF), University of Palermo, Viale delle Scienze, Ed. 5, 90128 Palermo, Italy

**Keywords:** MinION platform, Geminivirus, TYLCV recombinants, tomato, TYLCV-IMS54 recombinant

## Abstract

Tomato yellow leaf curl disease (TYLCD) is a major viral disease severely affecting tomato crops in the Mediterranean region, leading to reduced crop yield and significant economic losses. The disease is caused by monopartite begomoviruses belonging to the *Geminiviridae* family, primarily tomato yellow leaf curl Sardinia virus (TYLCSV) and tomato yellow leaf curl virus (TYLCV), which often co-infect tomato plants, promoting the emergence of recombinant viral genomes. To investigate the diversity and evolutionary dynamics of these viruses, symptomatic plants collected from agricultural sites in Sicily between 2020 and 2022, along with archived plant samples from 1994 to 1999, were analyzed. For each collection site, leaves from symptomatic plants were pooled to form representative samples. Total nucleic acids were extracted and subjected to rolling circle amplification to enrich circular viral genomes. The amplified products were sequenced using Oxford Nanopore Technologies (ONT) long-read sequencing to obtain full-length viral genomes. Bioinformatic analyses revealed that archived samples exclusively contained TYLCSV-related sequences, confirming its historical predominance in Sicilian agroecosystems. Recent samples, by contrast, no longer contained TYLCV or TYLCSV parental genomes but were dominated by TYLCV-derived recombinants such as TYLCV-IS141- and TYLCV-IS76-like variants, indicating a temporal shift in the structure of the viral population. Furthermore, a distinct group of newly emerged recombinants, provisionally referred to as TYLCV-IMS54, was identified in the most recent samples. Their genome comprises a TYLCV backbone, a 54-nucleotide segment from TYLCSV located downstream of the stem-loop region, and an 341-nucleotide region derived from TYLCV-Mild. These results demonstrate the importance of continuous viral population monitoring through ONT-based sequencing to detect emerging variants that may influence disease management strategies in tomato crops and highlight the central role of recombination in shaping begomovirus populations.

**IMPACT STATEMENT:** Tomato yellow leaf curl disease (TYLCD) is one of the most damaging viral diseases affecting tomato crops in the Mediterranean basin, yet the long-term dynamics of its causal agents and the role of recombination remain challenging due to the genome plasticity of these viruses. This study provides an updated and comprehensive picture of the begomovirus population structure in Sicily, a key agricultural region for tomato production, by analyzing both contemporary and historical plant samples. Through the application of ONT long-read sequencing combined with RCA and bioinformatic analyses, this research identified persistent recombinant genotypes including a distinct group of newly emerged recombinants, named TYLCV-IMS54.

These findings expand current knowledge on the genetic variability and evolutionary processes shaping begomovirus populations in Sicilian agroecosystems. The detection of recombinant genomes highlights the enduring role of recombination in begomovirus diversification. By integrating sequencing data with population and phylogenetic analysis, this work offers valuable insights into the epidemiology and management of TYLCD in regions heavily impacted or newly colonized by these viral pathogens. The study also underscores the importance of continuous molecular surveillance using ONT-based platforms to enable early detection of emerging recombinant variants, with significant implications for plant virology, crop protection and agricultural biosecurity strategies.

**DATA SUMMARY:** Raw reads are deposited in the Sequence Read Archive (SRA) of NCBI (https://www.ncbi.nlm.nih.gov/sra) with BioProjects ID PRJNA1226414 and PRJNA1273745; the recombinant TYLCV-IMS54 sequence is available in GenBank with Accession Number PQ873011. Parental TYLCV-strains used in this study could be retrieved in GenBank with the following Acc. Nos: DQ144621 for TYLCV, NC_003828 for TYLCSV, KJ913682 for TYLCV-Mild. Recombinant strains could be retrieved in GenBank with the following Acc. Nos.: LN846609 for TYLCV-IS76 and AF271234 for TYLCMaV.

The authors confirm that all supporting data, code and protocols have been provided within the article or through supplementary data files.

## INTRODUCTION

Tomato yellow leaf curl disease (TYLCD) is a major disease of tomato crops, causing significant economic losses worldwide. TYLCD is induced by viruses of the genus *Begomovirus* (family *Geminiviridae*) [1], one of the largest and most economically important group of plant-infecting viruses, transmitted in a persistent-circulative manner by the whitefly *Bemisia tabaci*. The rapid spread of TYLCD is exacerbated by the dynamic nature of the geminivirus genomes, which exhibit high mutation and recombination rates that enable them to rapidly adapt to new hosts and environmental conditions [2]. In Italy, two predominant TYLCD-associated species have been identified, tomato yellow leaf curl Sardinia virus (TYLCSV) (*Begomovirus solanumflavusardiniaense*), first reported in Sardinia in 1989 [3, 4], and tomato yellow leaf curl virus (TYLCV) (*Begomovirus coheni*), detected in Sicily in 2002 [5]. The co-existence of these two species resulted in the emergence of various recombinants, highlighting a high evolutionary flexibility [6]. The widespread cultivation of tomato varieties carrying resistance genes, particularly *Ty-1*, further shaped viral population dynamics. Indeed, while these resistance genes initially provided effective control of the disease, they exerted a selective pressure on the viral population, leading to the appearance of resistance-breaking variants [7]. Understanding the genetic diversity and adaptability of these viruses requires robust tools capable of capturing the complexity of their genomic evolution. To identify TYLCD-related isolates and recombinants, various diagnostic techniques are employed, each with specific advantages in sensitivity, specificity, and applicability, ranging from Restriction Fragment Length Polymorphism analysis followed by Polyacrylamide Gel Electrophoresis, and hybridization-based approaches [8, 9], to more advanced molecular techniques such as singleplex or multiplex Polymerase Chain Reaction (PCR) [10, 11], Loop-Mediated Isothermal Amplification (LAMP) [12], and the combination of LAMP with CRISPR-Cas [13]. Recent breakthroughs in sequencing technologies are continuously revolutionizing the study of viral diversity, evolution and recombinant detection [14]. In particular, the Oxford Nanopore Technologies (ONT) MinION platform has emerged as a powerful tool for the real-time identification of viral genomes [14]. Its capacity to generate long-read sequences facilitates precise detection of genetic variations, including mutations and recombination events, which are key drivers of viral evolution [15]. Furthermore, the portability and cost-effectiveness of ONT MinION make it a valuable tool for field-based applications and rapid diagnostics, enabling researchers to address challenges in TYLCD management with unprecedented efficiency [16].

In this study, using the combination of Rolling Circle Amplification (RCA) and ONT MinION platform, we provide a comprehensive overview of the genetic diversity and population dynamics of TYLCD-associated viruses in tomato and cucurbit plants across Sicily, a Southern Italian region where this disease causes significant agricultural losses. The evolutionary mechanisms underlying the adaptation of these viral populations are discussed.

## METHODS

### Sample collection and DNA extraction

During the years 2020-2022, five tomato (*Solanum lycopersicum*) fields showing TYLCD symptoms in the Ragusa province and one watermelon (*Citrullus lanatus*) field with atypical strong leaf mosaic symptoms in the Agrigento province were visited (Figure 1). From each field, young leaves from 20 randomly selected symptomatic plants were collected and pooled to create a single sample representing the whole field. Total nucleic acids were extracted from each sample (Table 1) using the previously described Dot-Blot method [17]. Nucleic acid concentration was quantified using a NanoDrop spectrophotometer (Thermo Scientific, Wilmington, USA). Additionally, DNA extracted from tomato and weed samples collected in the Ragusa province between 1994 and 1999 and archived at -20°C in our laboratory were used to investigate the evolutionary dynamics of TYLCD.

**Table 1.** List of samples analyzed by minION sequencing, consisting of pools of symptomatic leaves collected in Sicily in the years 1994 - 2022.

| Sampling year | Sample ID | MinION code | Location | Species/Cultivar | Genotype | Number of plants pooled in the sample |
| --- | --- | --- | --- | --- | --- | --- |
| 1994 | 051294_vv | D1K8_016 | RG | *Malva* spp.,<br>*Chenopodium album*,<br>*Potentilla* spp.,<br>*Portulaca* spp. and<br>other weeds | N.A. | 10 |
| 1999 | 130999_t_1 | D1K8_017 | RG | Tomato | U | 10 |
|  | 130999_t_2 | D1K8_018 | RG | Tomato | U | 10 |
| 2020 | 061020_t_2 | HPCT001 | Acate (RG) | Tomato/Creativo | T | 20 |
|  | 071020_t_3 | HPCT002 | Scicli (RG) | Tomato/Cikito | S | 20 |
| 2021 | 141021_t_4 | D1K8013 | Vittoria (RG) | Tomato/Cherry | T | 20 |
|  | 141021_t_5 | D1K8007 | Vittoria (RG) | Tomato/Datterino | U | 20 |
|  | 150721_w_1 | D1K8010 | Licata (AG) | Watermelon | U | 20 |
| 2022 | 141122_t_1 | D1K8009 | Vittoria (RG) | Tomato/Demetrio | T | 20 |

RG, Ragusa; AG, Agrigento; N.A., not applicable; U, unknown; T, tolerant; S, susceptible.

**Figure 1.**
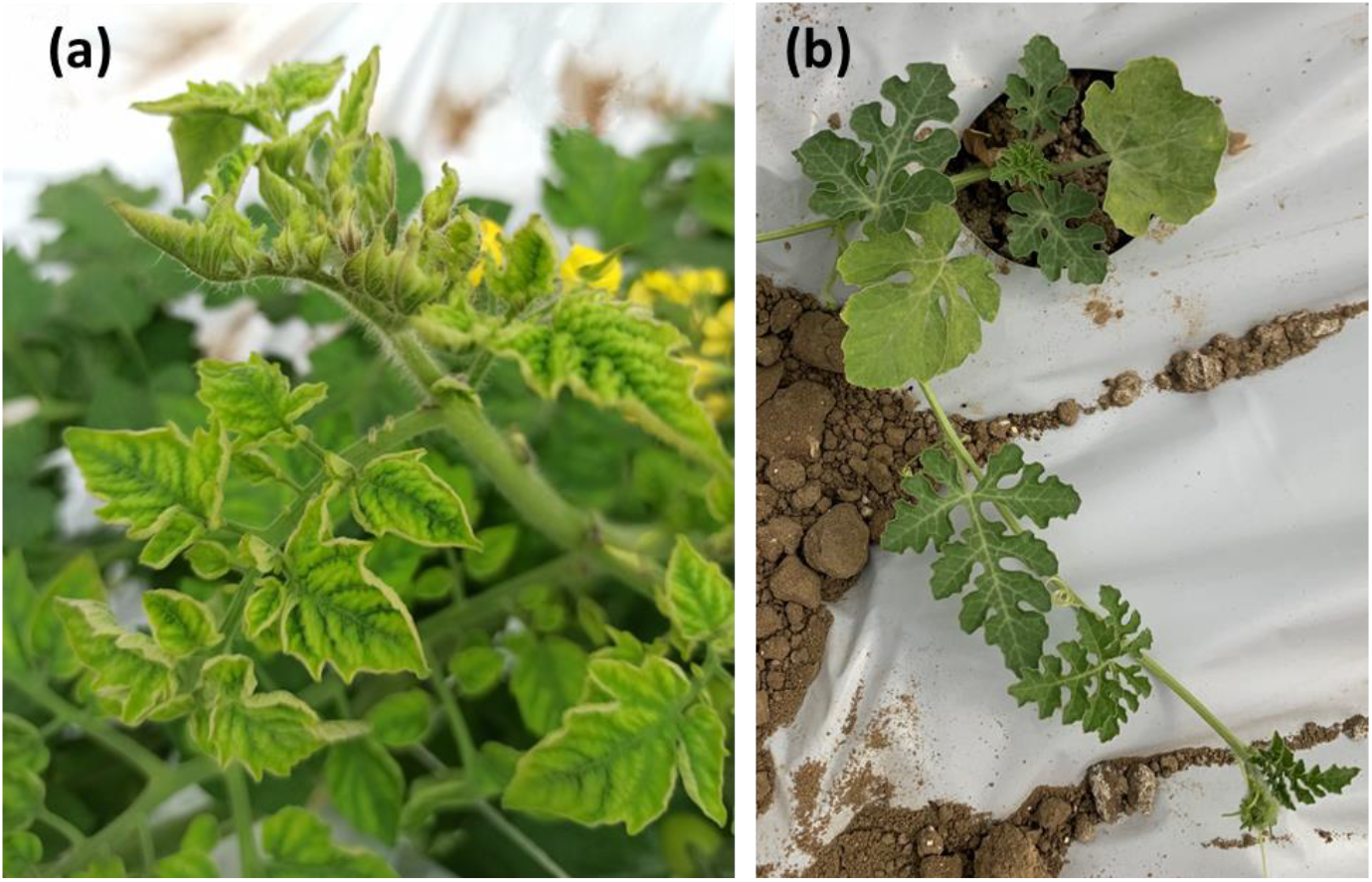
Geminiviral symptoms in plants tested in this work. (a) Tomato plant var. Demetrio (TYLCV tolerant) (sample 141122_t_1). (b) Watermelon plant showing chlorosis, mosaic and stunting (sample 150721_w_1).

### Rolling Circle Amplification

To enrich circular DNA molecules, Rolling Circle Amplification (RCA) was performed on total nucleic acids extracts using the TempliPhi™ 100 Amplification Kit (Cytiva 25-6400-10, Sigma Aldrich, St. Louis, MO, USA), running the reaction at 30°C for 30 hours. One-tenth volume of the RCA product was digested with either *BamH*I or *EcoR*I enzymes; linearized DNAs were resolved by electrophoresis on 1% agarose gels in 0.5X TBE buffer and visualized following staining with RedSafe™ (iNtRON Biotechnology, Korea), using undigested RCA products as control. The latter were precipitated with 3 volumes of ice-cold ethanol, and 0.1 volume of 3M sodium acetate and DNA was resuspended in 20 uL of sterile water and quantified by NanoDrop spectrophotometer.

### MinION sequencing and recombination analysis

Undigested RCA products were submitted to a specialized sequencing company (MicrobesNG, Birmingham, UK) for long-read sequencing on the Oxford Nanopore Technology (ONT) MinION platform, following manufacturer’s instruction. The MicrobesNG procedure included long-read libraries preparation using the Rapid Barcoding Kit 96 V14 (SQKRBK114.96) and sequencing with a R10.4.1 flowcell (FLO-MIN114) on GridION. Raw signal data were basecalled using the GridION deployment for Guppy (ont-guppy-for-gridion V 6.3.9) using model numberr1041_e82_400bps_hac_v4.2.0 with barcode trimming enabled. Reads under 200 base pairs (bp) were discarded. Raw long reads obtained from MicrobesNG were converted from FASTQ to FASTA format using Seqtk (v. 1.4-r130-dirty). Consensus sequences were assembled with TideHunter (v. 1.5.5) [18]. Sequences were subjected to a BLASTn search against a custom database comprising 16,957 *Geminiviridae* sequences downloaded from NCBI Virus (https://www.ncbi.nlm.nih.gov/labs/virus/vssi/#/, accessed on 19 May 2025, filters applied: GenBank and complete). For recombination analysis, geminivirus-related sequences longer than 2,770 nucleotides (nt) were checked with ORFfinder (https://www.ncbi.nlm.nih.gov/orffinder/) and multi-aligned with MUSCLE in MEGA11 (v. 11.0.13) [19] with putative parental TYLCV-related genomic sequences. Sequence Demarcation Tool (SDT v. 1.3) [20] was used to classify virus sequences based on sequence pairwise identity. Recombination events were subsequently identified using the Recombination Detection Program (RDP v. 4.101) [21]. Recombination events have been accepted only if identified by at least 5 different methods. Representative sequences of each recombination event were verified using BLASTn.

### End-point PCR analysis

Recombination breakpoints were validated by end-point PCR. Regions useful to design specific primers were selected following alignment of viral genomic sequences using ClustalW within BioEdit (v. 7.7.1) [22]. Primer specificity was assessed using Nucleotide-BLAST (https://www.ncbi.nlm.nih.gov, accessed on 7 June 2025) and the secondary structure was verified with the OligoAnalyzer Tool (https://eu.idtdna.com/calc/analyzer, accessed on 7 June 2025). The primer pair IMS54-2592-F (5’-GGAAVCGCTTAGGAGGAGCCAT-3’) and IMS54-138-R (5’-TTGCAAVACAAATTACTTGGGGA-3’) was designed to selectively amplify a region of about 330 nt. The assay included as control the following isolates DQ144621 for TYLCV, NC_003828 for TYLCSV, and KJ913682 for TYLCV-Mild, LN846609 for TYLCV-IS76, and AF271234 for TYLCMaV. DNAs from healthy tomato and cucurbit plants were used as negative controls. PCR was performed using Platinum™ II Taq Hot-Start DNA Polymerase kit (Invitrogen), following manufacturer’s instruction, with an annealing temperature of 58°C. PCR products were separated by electrophoresis as above. Selected PCR products were purified using the DNA Clean & Concentrator kit (Zymo Research) for Sanger sequencing analysis (BMR Genomics S.r.l. Padova, Italy).

### Phylogenetic analyses

To construct the phylogenetic tree, parental TYLCV-strains and recombinants from the Mediterranean area were downloaded from NCBI Virus (accessed on 23 May 2025) and multi-aligned with MUSCLE in MEGA11. A Neighbor-Joining (NJ) tree with 1000 bootstrap replicates was reconstructed with MEGA11. The sequence similarity level was estimated with BLASTn.

## RESULTS

### Identification of TYLCD-related genomic sequences

Using the ONT MinION platform combined with RCA, between 91 and 412 consensus sequences were obtained for each of the nine samples analyzed, as detailed in Table 2. To identify TYLCD-related sequences, reads longer than 2,770 nts were selected and used as input for a BLASTn search against the *Geminiviridae* custom database. Several TYLCD-related sequences (2 to 135) were identified. Sequences showing identity with TYLCD genomes were checked with ORFfinder to identify coding regions and only those with correct open reading frames (ORFs) (2 to 50) were retained for further analysis (Table 2).

**Table 2.** Oxford Nanopore Technology (ONT) MinION long reads statistics.

| Sample ID | Sample | Raw reads | Cleaned reads | Consensus sequences | Consensus sequences > 2,770 nts | TYLCD-related sequences | TYLCD-related sequences with correct ORFs |
| --- | --- | --- | --- | --- | --- | --- | --- |
| HPCT_001 | tomato | 4,130 | 2,094 | 119 | 35 | 35 | 2 |
| HPCT_002 | tomato | 3,144 | 1,597 | 91 | 22 | 22 | 2 |
| D1K8_007 | tomato | 2,280 | 1,152 | 202 | 80 | 78 | 10 |
| D1K8_009 | tomato | 4,011 | 2,029 | 219 | 33 | 28 | 2 |
| D1K8_013 | tomato | 2,371 | 1,348 | 218 | 39 | 26 | 5 |
| D1K8_010 | watermelon | 3,644 | 1,831 | 230 | 8 | 2 | 2 |
| D1K8_016 | weeds | 2,425 | 1,220 | 412 | 20 | 4 | 1 |
| D1K8_017 | tomato | 2,299 | 1,153 | 293 | 138 | 135 | 44 |
| D1K8_018 | tomato | 2,528 | 1,269 | 277 | 129 | 122 | 50 |

### Recombination analysis of samples collected in 2020-2022

Following alignment with the parental genomes, TYLCD-related sequences obtained from samples collected from 2020 to 2022 (hereafter referred to as r followed by a progressive number) showed levels of similarity to TYLCV or TYLCV-Mild higher than 94%, a value proposed as strain demarcation value for the *Begomovirus* genus by the *Geminiviridae* Study Group of the International Committee on Taxonomy of Viruses (ICTV) [23, 24]; based on this criterium, all the obtained sequences were assigned to TYLCV and none of them to TYLCSV (Figure 2).

**Figure 2.**
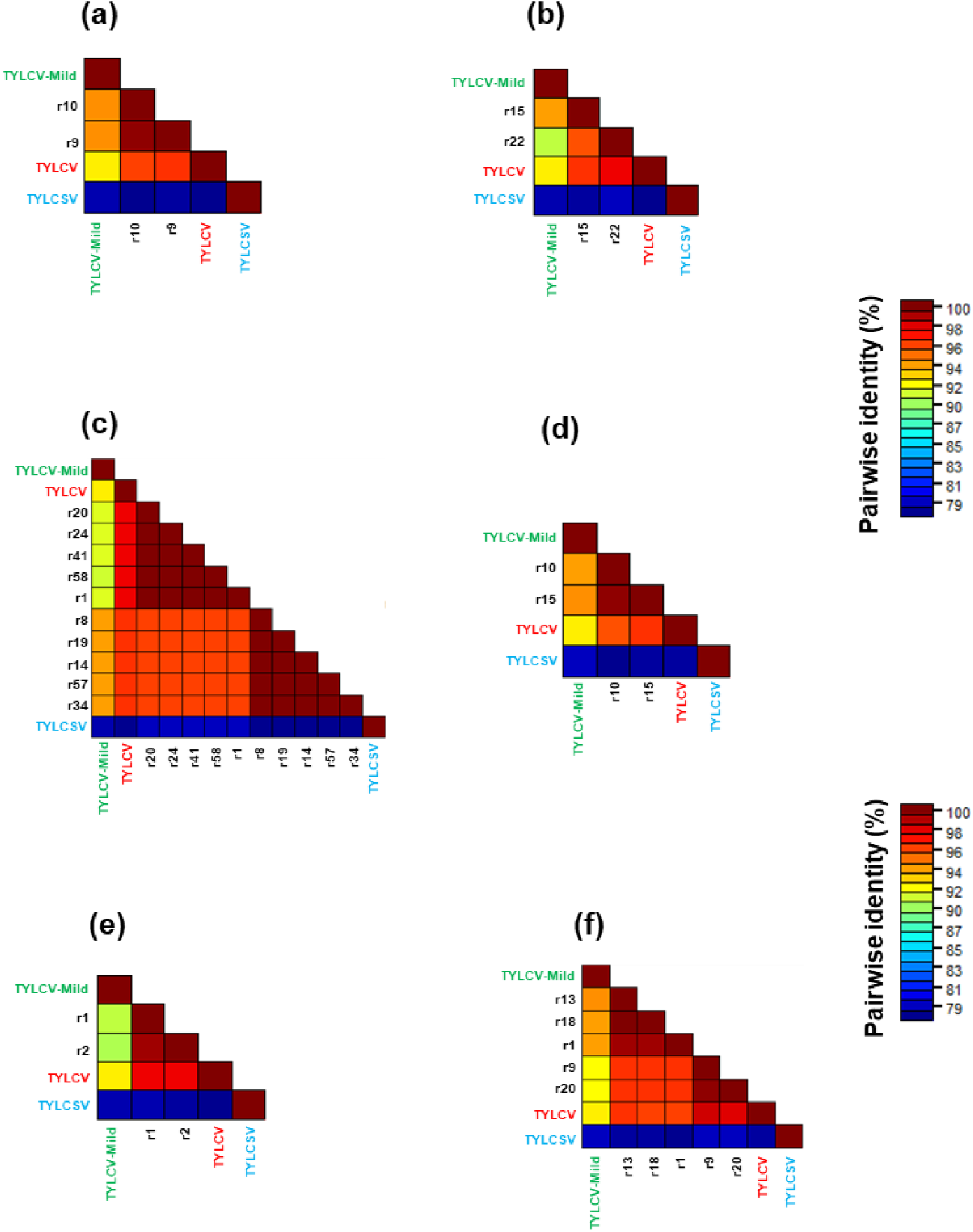
Pairwise identity matrices of TYLCD-related sequences obtained from each sample and the parental sequences from GenBank. Each matrix represents individual samples as follows: (a) HPCT_001, (b) HPCT_002, (c) D1K8_007, (d) D1K8_009, (e) D1K8_010, (f) D1K8_013. Recombinant (r) sequences were named with progressive numbers, within each sample. Parental genomes are written according to the following color code: TYLCV in red, TYLCV-Mild in green and TYLCSV in blue. Different clusters are identified at ≥ 94% identity, the defined value for strain demarcation of begomoviruses.

Pairwise identity analysis allowed us to identify the presence of clusters corresponding to different groups of TYLCD-related genotypes. Considering the complex scenario of the TYLCD-related sequences and the spread of recombinant genomes in the Mediterranean area, including the already known TYLCV-IS76, TYLCV-IS141, TYLCMaV, and TYLCAxV recombinants, we decided to more deeply characterize the sequences retrieved in our samples in order to identify possible recombination events and also define the presence or absence of parental strains, such as TYLCV and TYLCV-Mild. Following analyses with the RDP tool, up to four different kinds of recombination events in a single sample could be highlighted. Table 3 presents detailed results for each recombination event, including the associated P-values, the start and end points of the events, and the putative parental sequences.

**Table 3.**
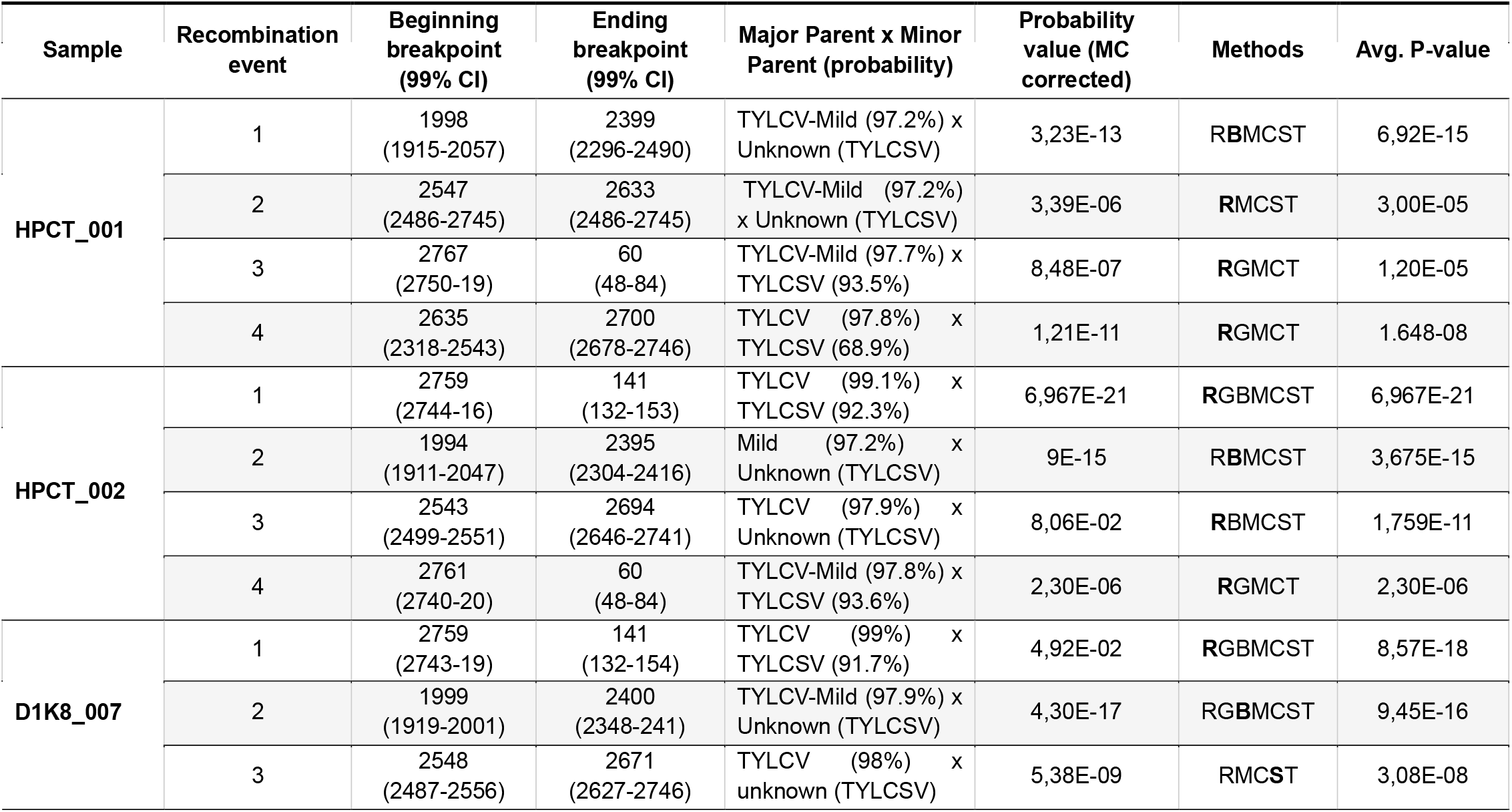
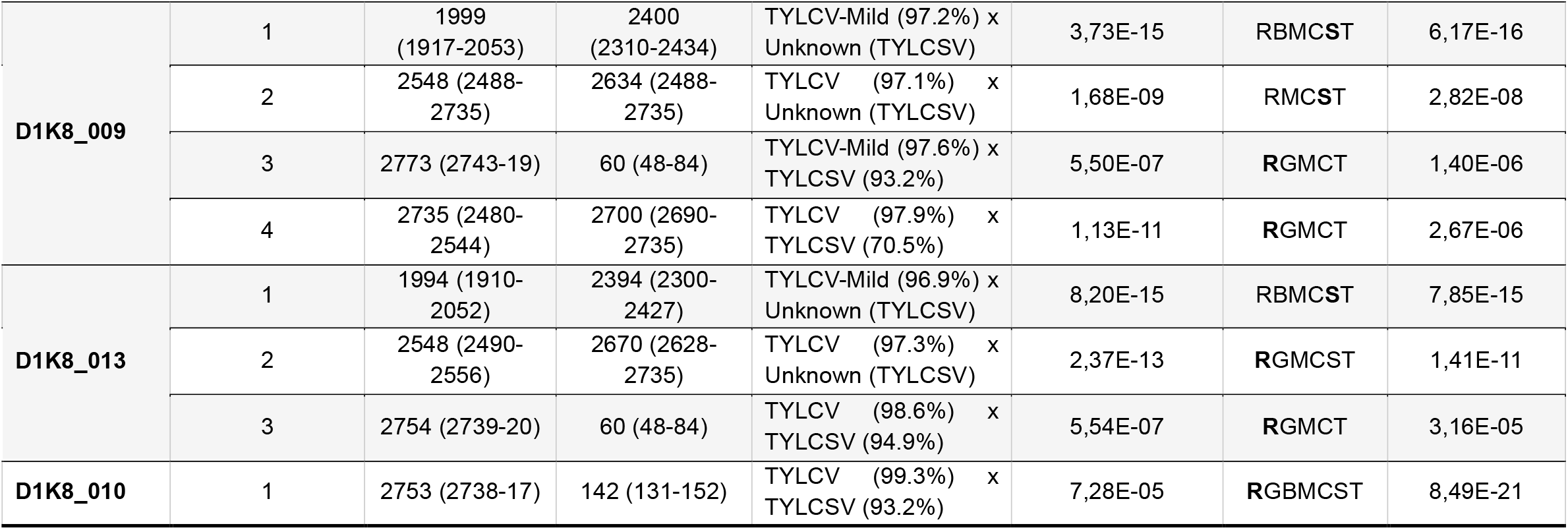
Recombination events identified by RDP in the TYLCD-related sequences. Beginning and ending breakpoints are indicated with confidence intervals (CI) at 99%. The probability value indicates the probability of observing a recombination signal, corrected for multiple testing, in that interval. When the minor parent is unknown, the inferred sequence is indicated in brackets. MC indicates Multiple Comparisons. Methods used to identify the recombination event are R = RDP; G = GENECONV; B = BootScan; M = MaxChi; C = Chimaera; S = SiScan; T = 3Seq; only methods supporting that event are listed. Average P-value is the lowest calculated and is referred to the method highlighted in bold.

Overall, this study highlighted the widespread presence of the already known TYLCV-IS141-like recombinant, detected in almost all the samples, and a limited presence of the TYLCV-IS76-like recombinant in tomato plants (sample D1K8_013) in the same geographical area. In the watermelon sample (D1K8_010), a TYLCV-IS141-like molecule was the only recombinant identified. Interestingly, in all the tomato samples a recombinant molecule putatively having TYLCV-Mild and TYLCSV as major and minor parents, respectively, was found, suggesting a high fitness of this viral genome.

Importantly, the parental genome sequences of TYLCV and TYLCSV, previously commonly found in Sicily [3-5], could not be detected in any of the samples here considered, supporting the above results obtained with SDT (Figure 2), indicating that fitter recombinant genomes had displaced the parental viruses over time.

### Recombination analysis of samples collected in 1994 and 1999

Thanks to the availability of new powerful sequencing techniques, such as ONT MinION pipeline, we decided to re-analyze old samples, collected in the same geographical areas, where TYLCSV sequences only were reported [8]. The pairwise identity analysis highlighted the existence of two different clusters of sequences in each of the two tomato samples D1K8_017 and D1K8_018 (Figure 4). These clusters represent two different groups of TYLCSV-related isolates previously identified in Sicily in 1995 (Z28390) and in 2004 (GU951759). Similarly, the unique sequence obtained from the mixed weed sample showed a high level of identity (99.64%) with the sequence of the Sicilian isolate Z28390.

**Figure 3.**
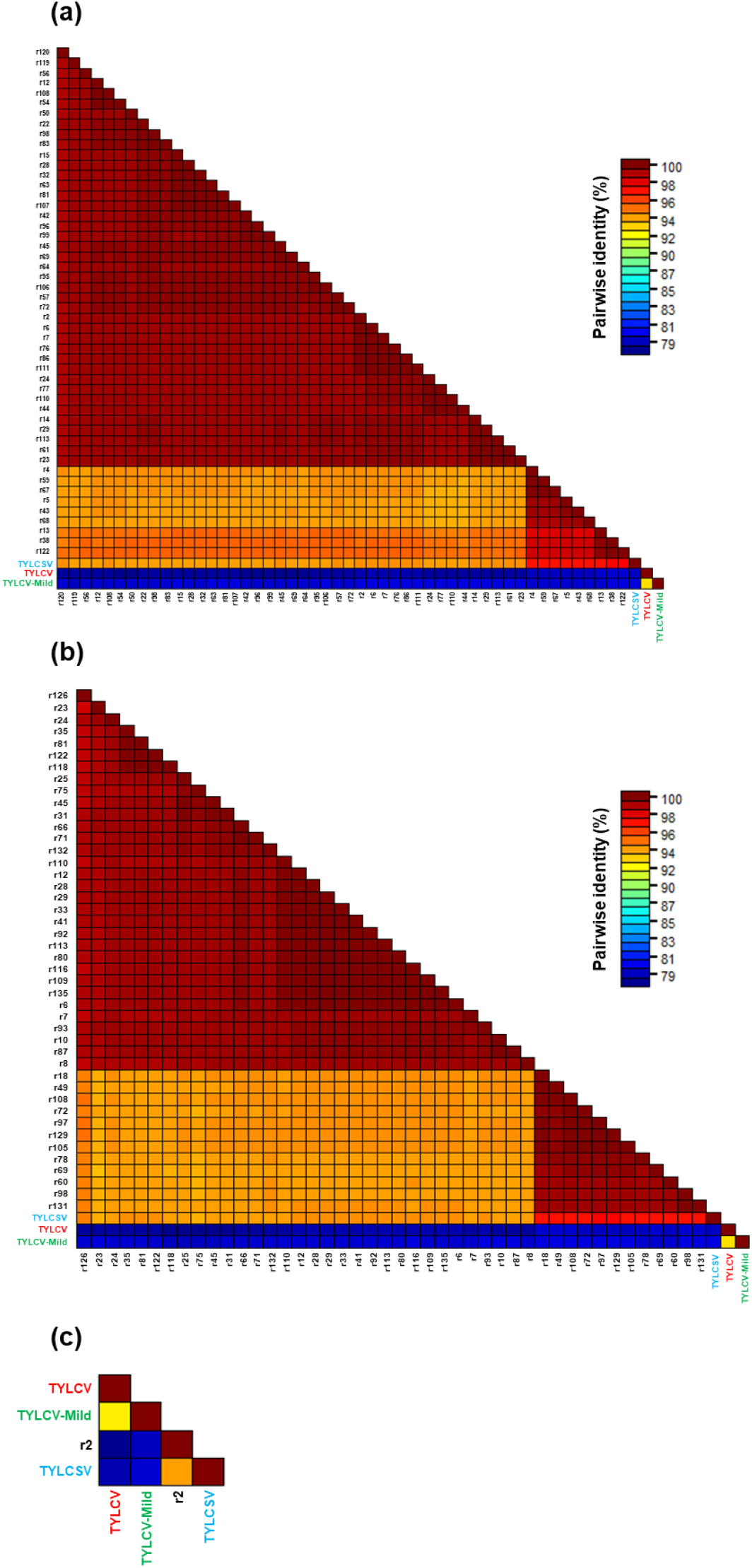
Pairwise identity matrices of TYLCD-related sequences obtained from each sample and the parental sequences from GenBank. Each matrix represents individual samples as follows: a) D1K8_017; b) D1K8_018 and c) D1K8_016. Recombinant (r) sequences were named with progressive numbers, within each sample. Parental genomes are written with the following color code: TYLCV in red, TYLCV-Mild in green and TYLCSV in blue. Different clusters are identified at ≥ 94% identity, the defined value for strain demarcation of begomoviruses.

**Figure 4.**
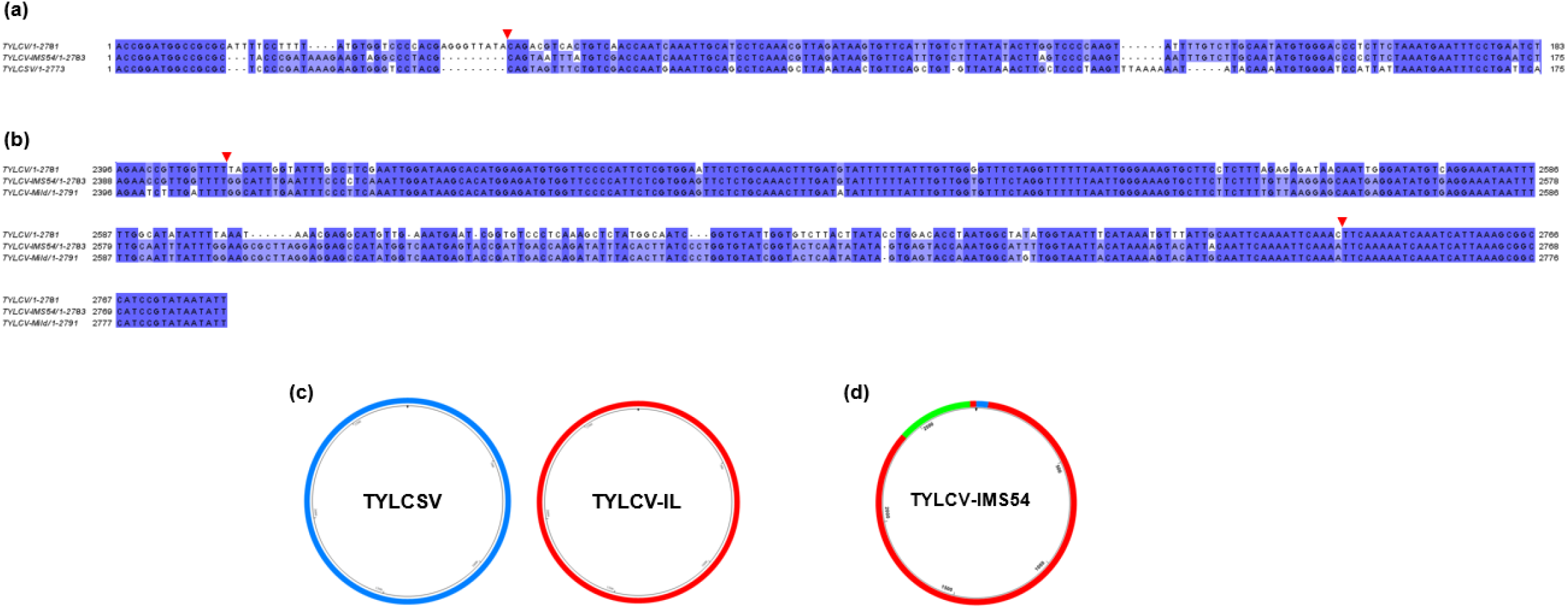
Features of the TYLCV-IMS54 recombinant. a) DNA alignment of nucleotides 1-175 of TYLCV (DQ144621), TYLCSV (NC_003828) and TYLCV-ISM54 (PQ873011); b) DNA alignment of nucleotides 2388-2783 of TYLCV (DQ144621), TYLCV-Mild (KJ913682) and TYLCV-ISM54 (PQ873011). In both a) and b), the red arrow indicates the recombination point. c - d) Schematic representation of the genomes of parental (c) and recombinant (d) isolates relevant to this study. TYLCV is represented in red, TYLCSV in blue and the sequence portions characterizing TYLCV-Mild are in green.

The results confirmed the sole presence of TYLCSV in that time frame (Figure 4), supporting the hypothesis that the spread of TYLCV and consequently the onset of TYLCSV/TYLCV recombinants occurred after 1999, at least in this intensive tomato farming area.

### Characterization of the TYLCV-Mild/TYLCSV recombinant

To investigate the nature of the recombinant molecule identified in the tomato samples from the years 2022-2022, putatively having TYLCV-Mild and TYLCSV as parentals (Table 3), an in-depth characterization was performed. The RDP analysis highlighted the insertion of a fragment of about 54 nts showing 98-100% identity with TYLCSV immediately on the right side of the stem-loop, while the remaining sequence matched with TYLCV-Mild. However, a more detailed analysis conducted with BLASTn on the genomic portions differentiating TYLCV-Mild from TYLCV indicated that the region between 1998-2399 nts matched with TYLCV, with an identity of 99%, while the region between 2415-2755 nts matched with TYLCV-Mild, with an identity of 100%. These results show that the recombinant molecule has a TYLCV backbone with a portion of TYLCV-Mild and one of TYLCSV. Its tripartite nature led us to name it TYLCV-IMS54; the corresponding sequence derived from sample D1K8_009 was deposited in GenBank with the Acc. No. PQ873011.

Recombination events involving TYLCV-Mild and TYLCSV were previously reported on common bean in Almeria (Spain), consisting in the TYLCMaV genome [25]. However, differently from TYLCMaV that included a TYLCSV portion of about 1,130 nts, the recombinant TYLCV-IMS54 is characterized by an extremely reduced TYLCSV contribution (Figure 4).

PCR analysis with dedicated primers followed by Sanger sequencing confirmed the presence of TYLCV-IMS54 in the original tomato samples (data not shown).

### Phylogenetic analyses

Phylogenetic analysis showed that the TYLCV-IS141-like recombinants found in this study cluster together with TYLCV-IS141 recombinant types of Italian origin, but distantly from the unique TYLCV-IS141 isolated in France (MG489967). Similarly, our TYLCV-IS76-like sequence clusters with the TYLCV isolate 8-4/2004 and a TYLCV from Tunisia, rather than with the sequences annotated as TYLCV-IS76 from Morocco and Spain (Figure 5). These findings likely indicate the existence of independent recombination events in different geographical areas.

**Figure 5.**
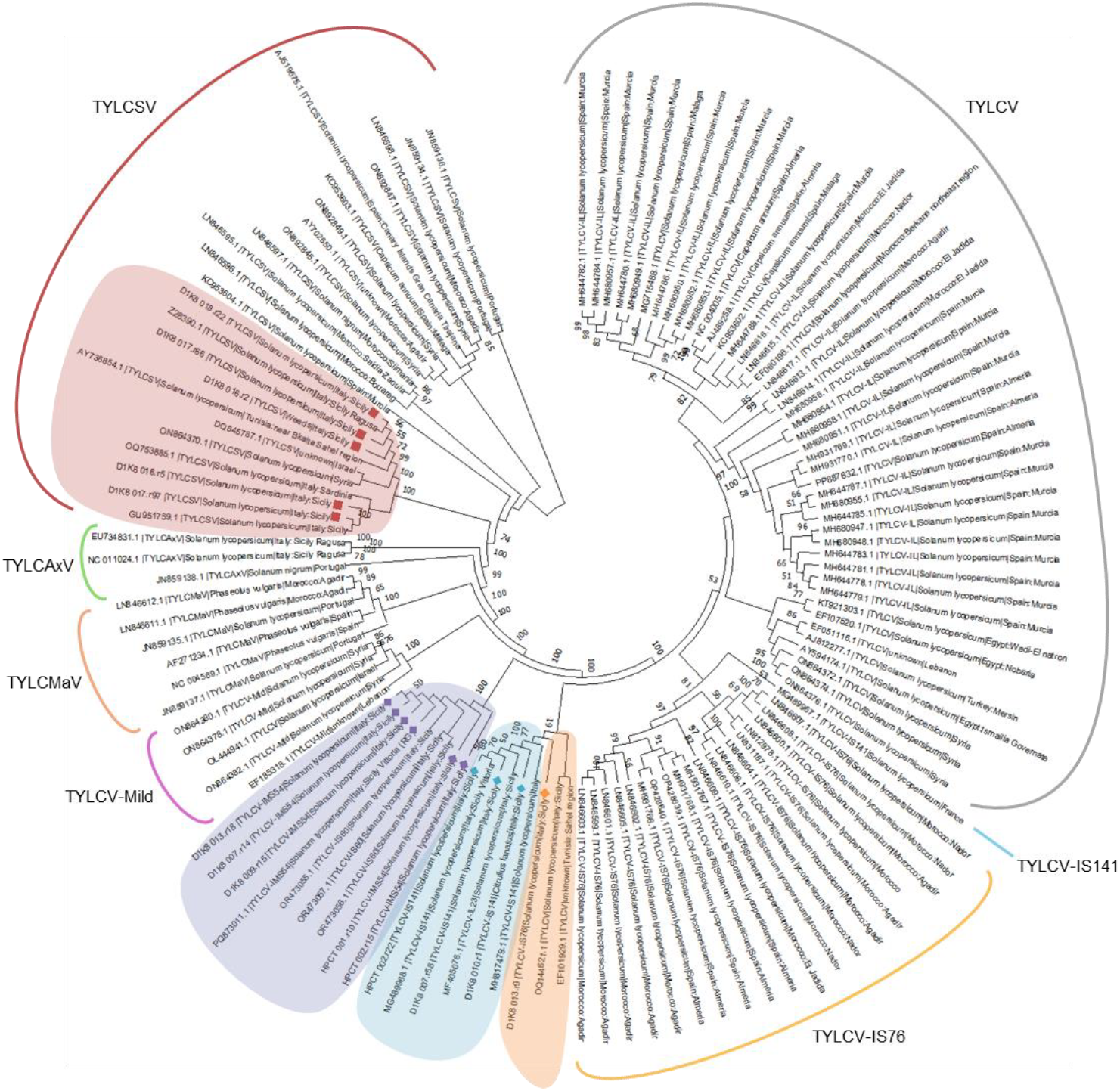
Neighbor-Joining (NJ) phylogenetic tree of TYLCV/TYLCSV recombinants (highlighted by colored diamonds) and parentals (highlighted by colored squares) obtained in this work, challenged with full-length TYLCD-related sequences from the Mediterranean area, downloaded from NCBI Virus. TYLCV-IS76 = orange; TYLCV-IS141 = turquoise; TYLCVIMS54 = purple; TYLCSV = red.

The recombinant TYLCV-IMS54 sequences cluster with the TYLCV-IS60 recombinants reported in Sicilian tomato samples in 2024 [11] and recently deposited in GenBank database (OR473055, OR473056, OR473057) (Figure 5). Indeed, our TYLCV-IMS54 type sequence (PQ873011) shows more than 99.7% identity with the three TYLCV-IS60 sequences. In light of this high similarity, a recombination analysis including TYLCV-Mild as putative parental, was performed on the three TYLCV-IS60 sequences mentioned above. The results highlighted that the TYLCV-IMS54/TYLCV-IS60-like sequences represent a new group of recombinants whose genome is composed of the three parentals TYLCV (major parent), TYLCV-Mild, and TYLCSV. Again, the existence of a single clade including all these sequences highlights their independent origin in Sicily (Figure 5).

## DISCUSSION

MinION sequencing following circular DNA molecules enrichment has emerged as a powerful and affordable tool for characterizing the geminivirus population infecting crops, addressing potential limitations of second-generation sequencing methods [15, 16]. Indeed, although the base-calling accuracy of Nanopore technologies still does not match that of second-generation sequencing platforms, their main advantage lies in the ability to achieve greater assembly completeness, owing to longer read lengths that are well suited to the size of geminiviral genomes [15]. In this study, we took advantage of the MinION sequencing technology to investigate the population dynamics of invasive TYLCD-related viruses in Sicily, a region of intensive tomato cultivation.

First of all, we highlighted a shift in the TYLCD*-*associated viral landscape, with the complete displacement of parental viral genomes over the years in favor of TYLCV recombinants with an enhanced fitness such as TYLCV-IS141- and TYLCV-IS76-like recombinants [32]. Interestingly, the detection of the TYLCV-IMS54 / TYLCV-IS60 recombinant in Sicily from as early as 2016 [11] through to our 2022 survey suggests that this variant also exhibits a high level of fitness, raising concerns about its potential impact on crop production. This recombinant is characterized by the introgression of a portion of 50-60 nts of TYLCSV in the intergenic region, immediately after the stem loop and by a portion of 341 nts originating from TYLCV-Mild on the left side of the genome. The presence of the small portion of TYLCSV seems to be crucial for the high fitness, thus reinforcing the conclusions reported by [32]. However, we cannot exclude that the TYLCV-Mild insertion could also have a role in further increasing the fitness and therefore further investigations are needed.

TYLCV-Mild and TYLCSV were reported to infect watermelon and other cucurbits in Jordan, producing leaf yellowing, curling and mottling symptoms [26] and TYLCV was detected in watermelon in Tunisia [27], in squash in Cuba [28], in cucumber in Kuwait [29], and in *Cucurbita maxima* in Japan [30]. To our knowledge, TYLCD-related viruses were never detected in Europe/Italy on cucurbits, and this is the first report of a TYLCV-IS141-like recombinant in Italy. The lack of PCR amplification with *B. tabaci*-specific primers [31] from our sample (data not shown) allowed us to exclude contamination by infected whiteflies. Our findings imply that TYLCV-IS141-like recombinants can infect cucurbit species, further confirming its superior fitness compared to parental genomes.

## CONCLUSION

In conclusion, our study confirmed the effectiveness of RCA combined with ONT technology as a robust and cost-effective approach for characterizing geminivirus populations in the field, enabling real-time monitoring of their spread. Through this technology, we documented a dynamic shift in the TYLCD-associated viral population in Sicily, with the displacement of parental viral strains by highly fit TYLCV recombinants such as TYLCV-IS141-, TYLCV-IS76- and TYLCV-IMS54/IS60-like variants. Our findings underline the adaptive advantage of recombination and emphasize the importance of continuous monitoring in order to develop effective management strategies and mitigate future risks for cultivation.

## Author statements

### Author contributions

E.N. and A.M.V. conceived and designed the study. E.N., A.M.V., L.M., S.D. and G.P.A. collected the samples and oversaw sample storage logistics. S.B., F.F. and A.M.V. performed wet-lab analyses. S.R., S.B. and L.M. conducted *in silico* analyses. A.M.V. and G.P.A. acquired the fundings. A.M.V., E.N., L.M., S.B. and S.R. co-wrote the manuscript, and all authors approved the final version of the manuscript.

## Acknowledgements

The authors wish to thank Daniele Marian and Slavica Matić for technical support during sample collection and Camilla Sacco Botto and Alessandro Aimone Catti for assistance with figure preparation.

